# Promera: a unified model for biomolecular structure prediction, filtering, and design

**DOI:** 10.64898/2026.06.07.729267

**Authors:** Bowen Jing, Mihir Bafna, Daniel Diaz, Adam Klivans, Bonnie Berger

## Abstract

Generative models have become staple tools for modeling and designing biomolecular structures. However, although these tools have improved in structural prediction accuracy, their ability to filter designed binders—an essential use case—remains insufficient; whereas design methods have focused more on unconstrained binder generation rather than capabilities enabled by controllable design. We introduce **Promera**, a unified generative model that combines all-atom structure prediction with improved filtering and controllable design. We find that Promera’s confidence metrics are more accurate for filtering binders from non-binders for both miniproteins and nanobodies, while its co-folding performance surpasses popular open-source models (OpenFold3-p2, Boltz-2) on therapeutically relevant categories. As a design model, Promera generates binders by predicting masked protein sequences with optional epitope, paratope, and template constraints. Remarkably, our nanobody designs match the *in silico* success rates from backprop-based techniques (mBER) when evaluated under co-folding confidence filters. We further provide two *in silico* demonstrations of the the versatile capabilities of our design method: epitope targeting of the Andes hantavirus glycoprotein with VHHs and active state stabilization of the *β*_2_ andrenergic GPCR. We conclude by proposing a scaling law for co-folding models, suggesting a path for further performance improvement.

## Introduction

Generative models have significantly advanced our ability to computationally model and design biomolecular structures. A growing ecosystem of models, both open-source and commercial, aspires to transform drug discovery, protein engineering, and basic biological research from a painstaking experimental process to a fast computationally-guided one. Broadly speaking, this ecosystem comprises two modeling layers: (1) co-folding models [1, 32, 35, 29, 12] for predicting the structure of biomolecules; and (2) a diverse array of design protocols [24, 10, 21, 28, 11] or generative models [27, 7, 13] for designing binders to desired targets. Advances in these layers have also engendered significant commercial interest in generative modeling for drug discovery (e.g. *de novo* antibody design) [8].

However, current methods built on the open-source ecosystem have two key limitations. First, despite advances in structural accuracy, commonly used co-folding models exhibit insufficient ability to discriminate binders from nonbinders [26]. This shortcoming hobbles all generative design workflows, which rely heavily on co-folding confidence metrics to select variants for experimental validation from thousands of model samples. Second, open-source models have been largely developed for unconstrained binder design tasks, with the aim of substituting for traditional screening campaigns, rather than design settings where screening is more difficult in the first place, such as epitope or functional specificity. Overcoming these limitations could enable more reliable and routine generative design of functionally targeted binders.

Here, we present Promera, a unified model for co-folding, binder filtering, and design, that advances the open source ecosystem on these dimensions. As a co-folding model, we show Promera exceeds Boltz-2 and OpenFold3-p2 in protein-protein, antibody-antigen, and protein-ligand interaction categories. Unlike with previous models, we benchmarked and optimized Promera confidence metrics for binder filtering in both minibinder and antibody design settings. In a retrospective analysis of Adaptyv’s well-known Nipah virus protein design competition, our confidence metrics enrich binders two-fold over metrics from Boltz-2, AF-Multimer, and Protenix-v1. We further developed an *interface contact score* (iCS) for filtering antibody-antigen interfaces. When benchmarked on shuffled nanobody-antigen pairs, our iCS exceeds existing confidence metrics to attain up to 20-fold enrichment of binding pairs @ 10% recall. These results position Promera as the highest-signal model available for use in open-source filtering pipelines.

Promera is a versatile and controllable binder design model. Leveraging the observation that co-folding models generate structured backbones for masked residues [11], we incorporate masking as an integral part of our training pipeline to allow the model to be used for design. We additionally provide epitope, paratope, and template conditioning, so that binder backbones can be steered towards a target interface or receptor geometry. We demonstrate that Promera significantly exceeds the *in silico* nabobody design success rates of BoltzGen [27] and, remarkably, matches or exceeds mBER [28], a hallucination (backprop-based) BindCraft-style method commonly thought to be superior to generative methods [6]. We provide two *in silico* case studies to showcase Promera’s ability to tackle highly challenging, but therapeutically relevant design settings: (1) targeting several distinct epitopes of a viral protein; and (2) stabilizing the active conformation of a G-protein coupled receptor (GPCR). Through these demonstrations, we seek to democratize nominal design capabilities previously only accessible through commercial protein design models [30, 4, 5].

We conclude by proposing a scaling law for biomolecular structure prediction by aggregating our performance and training budget with those from other models. Our scaling law suggests that significantly enhanced co-folding performance—up to AlphaFold3 level and beyond—can be attained with increasing consumption of data and compute during model training. Although similar trends have been documented for protein language models in the context of variant effect prediction [3], our results are the first in the context of structural modeling and will have significant implications for future model development.

## Results

### Overview of model architecture and training

Promera takes as input a set of molecular identities (e.g., protein sequences, RNA/DNA sequences, or small molecules), with to-be-designed residues masked, and outputs an atomic structure of the interacting complex. Our model architecture is generally based on that of AlphaFold3 [1], including pairformer, diffusion, and confidence modules. We make modifications to this architecture to allow for paratope, epitope, and structural inputs in binder design, and to fix a potential ambiguity in small molecule featurization. These modifications are described in more detail in Appendix A. We train Promera on the PDB (2023-12-31 cut-off) as well as a protein monomer self-distillation set curated from Mgnify and AlphaFoldDB. To align the model training with backbone generation for design, we replace a fraction of standard amino acids with `UNK` tokens and remove their sidechain atoms in 5% of batches. Similarly, we provide epitope conditioning and partial distogram conditioning each in 5% of batches. Validation curves and additional details are provided in Appendix A, Figure 10.

### Promera co-folding accuracy matches or exceeds popular open-source models

We first sought to assess the co-folding performance of Promera on PDB complexes as compared to three recently released open-source co-folding models: OpenFold3-p2 (OF3), Boltz-2, and Protenix-v1 [25, 34, 33]. We tested each model on a benchmark set of 1605 PDB complexes released in 2025, comprised of low-redundancy interfaces with low homology to prior complexes (details in Appendix B). No complex in the 2025 PDB benchmark was seen during training or validation of any of the four models. We find Promera broadly outperforms OF3 and Boltz-2 across molecular interaction categories, in terms of both LDDTs and success rates (Figure 1). Promera also matches Protenix-v1 in protein-ligand accuracy, although slightly trails it in protein-protein accuracy. The comparative advantage on protein-ligand accuracy is potentially a result of our improved ligand featurization scheme (Appendix A).

**Figure 1.**
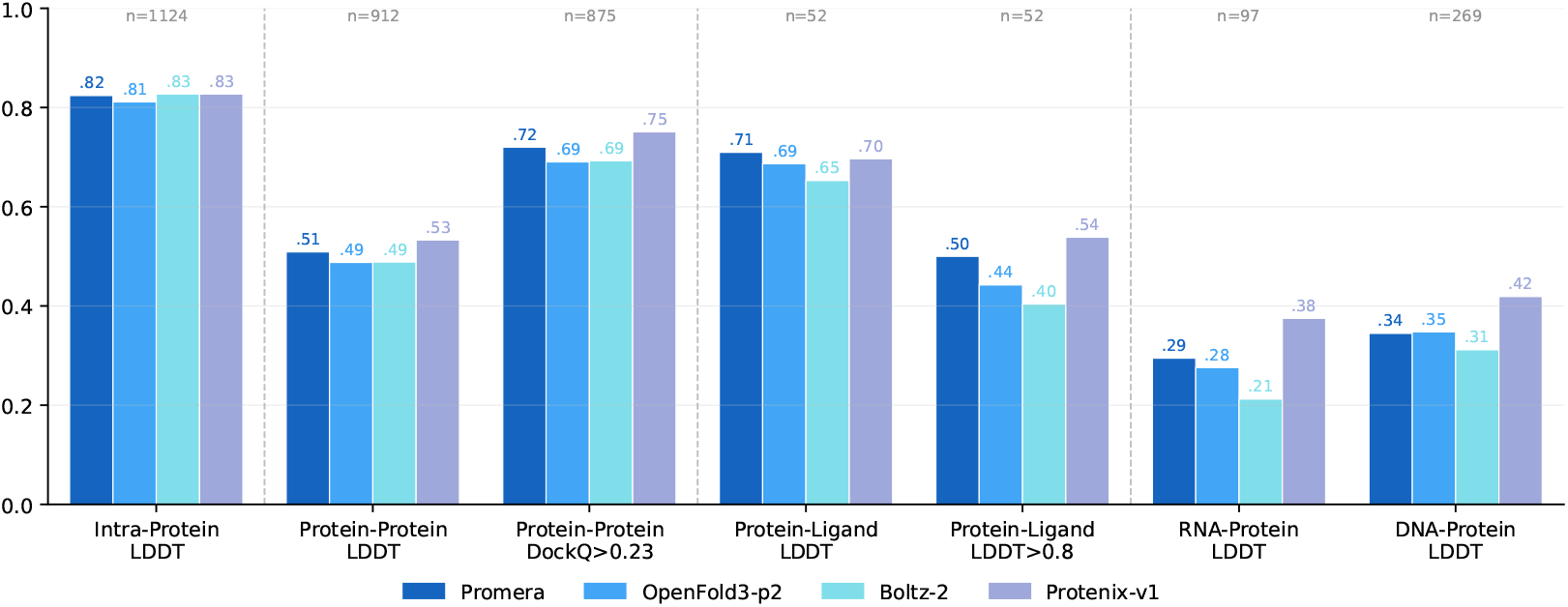
Co-folding evaluations on unseen PDB complexes. released in 2025. All values are reported as means across clusters, where each cluster may contain multiple chains or interfaces. Categories are annotated with the number of clusters. Detailed evaluation protocols are described in Appendix B. LDDT: Local Distance Difference Test.

We next sought to assess the performance of Promera on antibody-antigen co-folding by curating a test set from SabDab [15] entries released in 2025, deduplicated by clustering the antigen sequences at 50% similarity. This set comprises 221 complexes (146 Fabs, 6 scFVs and 69 VHHs). We show that Promera similarly outperforms OF3 and Boltz-2 on antibody-antigen co-folding (Figure 12). In particular, as the number of inference seeds increase, we observe a steady improvement in co-folding success rates using the top-ranked seed (e.g., from 20% to 27% at DockQ*>*0.49). This inference-scaling behavior, initially described in AlphaFold3, was noticeably absent from the first generation of open-source co-folding models (Boltz, Chai) [34]. Nevertheless, the oracle success rates of all methods exceed the ranked success rates, indicating substantial underlying sample diversity, but relatively poor selection of the best prediction.

We also document inference scaling behavior for protein-ligand co-folding. We reanalyzed model predictions for protein-ligand interfaces as a function of sample count in Figure 13. Using the best model prediction, the Promera protein-ligand success rate (LDDT>0.8) improves from 46% with one sample to 65% with 25 samples. The best-ranked prediction exhibits somewhat less improvement, from 46% to 50%. Nevertheless, extrapolating the observed trend suggests that scaling beyond the standard five seeds could improve protein-ligand accuracy. We leave this as a high-priority direction for further investigation.

### Promera confidence scores enrich binders over nonbinders

A major use case of structure prediction models is for filtering sequences in *de novo* protein design. Indeed, most recently published generative design protocols rely on (co-)folding confidence metrics to select designs for experimental validation out of a larger number (sometimes thousands) of samples. To date, however, there has been no systematic analysis of the ability of modern co-folding models to filter binders from nonbinders, especially in a *de novo* design setting. We address this gap by performing a systematic analysis of Promera and other top methods in both minibinder and nanobody settings (summarized in Figure 2).

**Figure 2.**
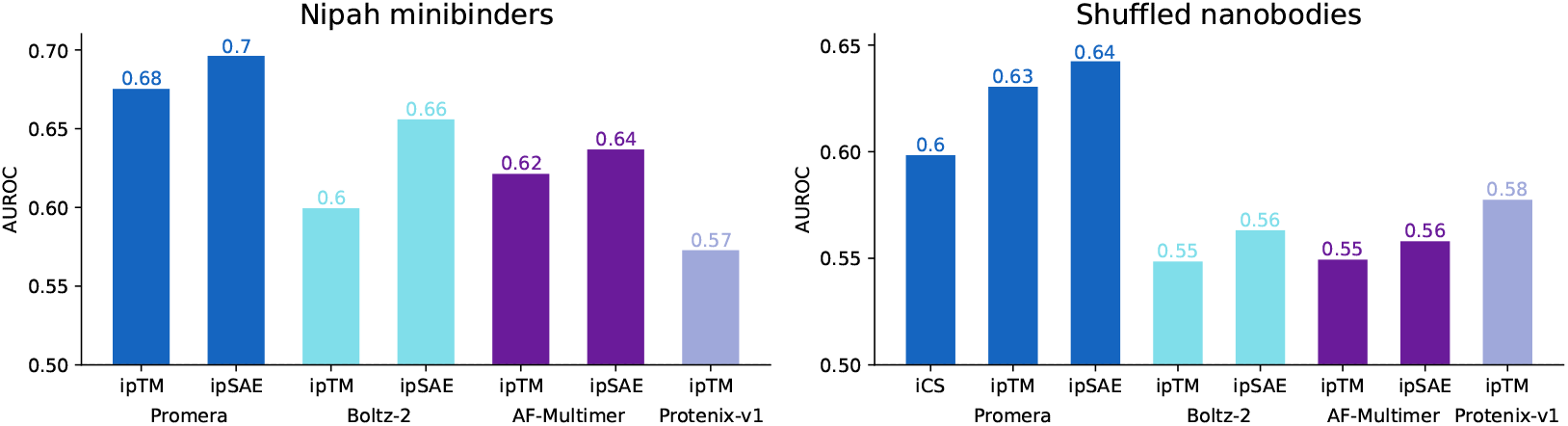
Binder discrimination ability of various confidence metrics as measured by AUROC on the Nipah minibinder dataset (**left**) and shuffled nanobody-antigen pairs (**right**). Promera confidence metrics have greater classification power than commonly used ones from Boltz-2, AF-Multimer, or Protenix-v1. ipTM: interface predicted TMscore; ipSAE: interaction prediction score from aligned errors.

#### Enrichment of minibinders from a recent design competition

To evaluate the ability of Promera and other methods to filter minibinders, we collected the sequences submitted to the 2025 Adaptyv Bio Nipah virus protein design competition. After excluding designs labeled as scFVs and VHHs, this dataset comprises 94 binders and 822 nonbinders, for 916 total minibinders. We then co-fold each binder sequence with the protein target sequence with each of Promera, Protenix-v1 [33], AlphaFold-Multimer (via ColabFold [22]), and Boltz-2 [25]. We compute the ipTM from each method; we also compute the ipSAE [16] for all methods except Protenix-v1 (which does not save full PAE matrices).

Promera ipSAE provides the greatest discimination power of binders versus nonbinders, with an AUROC of 0.70 compared to 0.66 for the next-best method (Figure 2, left). To provide a more fine-grained characterization of each confidence metrics, we then calculated the enrichment factor of binders over nonbinders as a function of score threshold (Figure 3, bottom left). The enrichment generally increases with higher score thresholds, at the cost of reduced recall of positive binders (Figure 3, bottom right). Promera ipSAE attains a higher maximum enrichment score than all other confidence metrics, attaining 5x at ≈10% recall, compared to 2x-3x for other methods. We emphasize that all 922 sequences were deemed to be likely binders by submitters and Adaptyv organizers, yet only 10% were successful. Notably, filtering these submissions with Promera to ≈ 20 samples could have led to a theoretical ≈ 50% success rate, a dramatic improvement.

**Figure 3.**
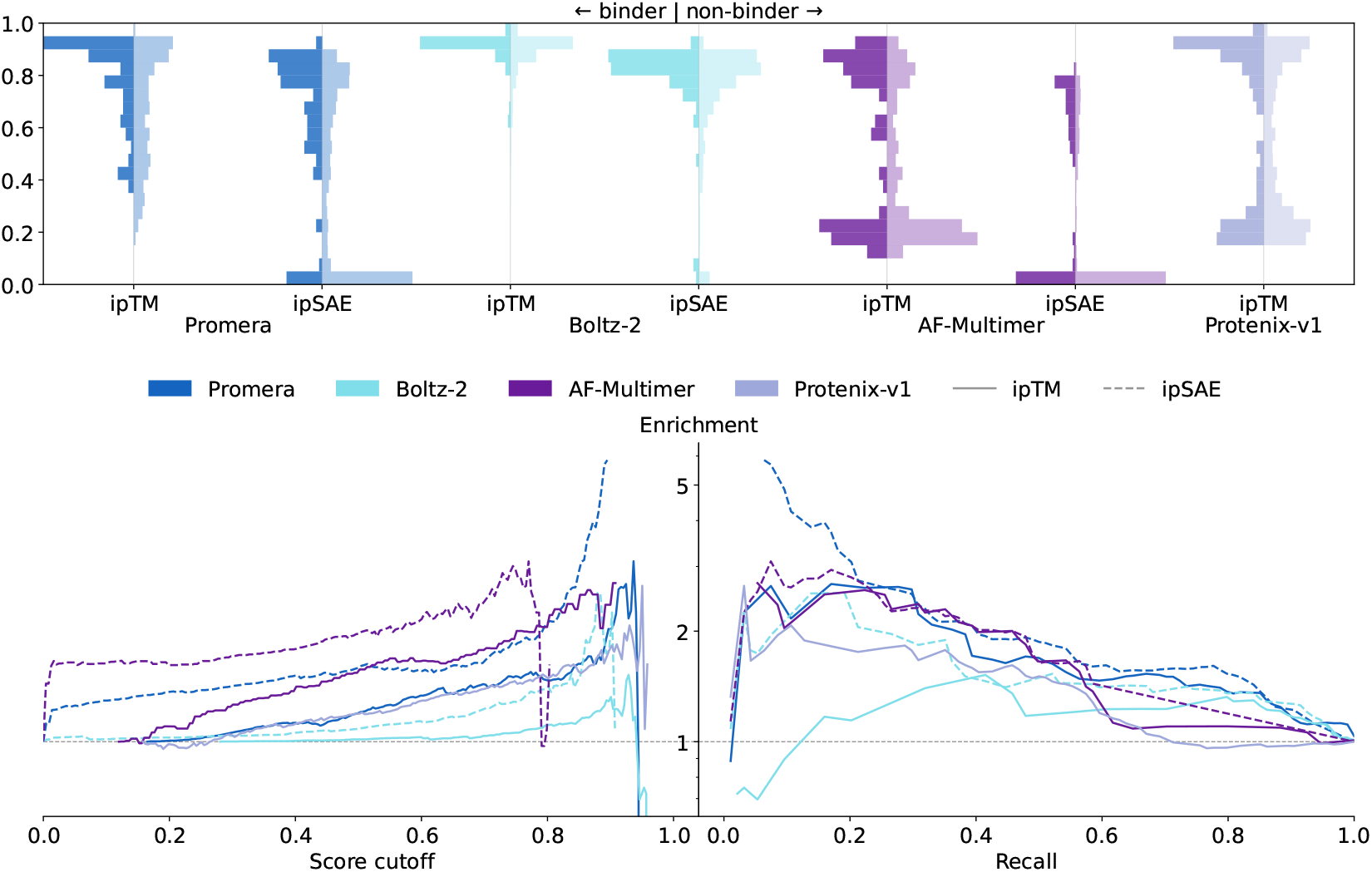
Discrimination of binders in the Nipah virus protein design competition with co-folding confidence metrics. (**Top**) Histograms of each score for binders and non-binders. An ideal scoring function would show a clear separation between binder and non-binder distributions. (**Bottom left**) Enrichment factors computed with each scoring method, as a function of score cutoff. Note that the curves are not directly comparable since the scores are not mutually calibrated. (**Bottom right**) Enrichment-recall tradeoff curves for each scoring function. For a given false negative budget (discarding true binders), the enrichment factor gives the expected improvement in hit rate over the baseline rate. Promera ipSAE attains higher enrichment than any other confidence score. ipTM (solid line): interface predicted TMscore; ipSAE (dashed line): interaction prediction score from aligned errors.

Since Boltz-2 ipSAE was used to select sequences for validation in the Nipah virus protein design competition, it may be at an artificial disadvantage. We therefore repeated a similar analysis using the older EGFR protein design competition (Appendix C), for which AF-Multimer ipTM was used to select designs.

#### High enrichment of nanobody-antigen interfaces from shuffled pairs

Antibody design forms a major subcategory of generative design settings and is of significant therapeutic relevance. Indeed, filtering designed antibodies likely comprises the plurality of co-folding production runs in industry, despite poor characterization of this capability in existing models. Here, we assessed the ability of Promera and other co-folding methods to filter binding from nonbinding antibody-antigen pairs. To do so, we extract all VHH-antigen interfaces from the 2025 SabDab antibody test set, resulting in 69 interfaces. Inspired by the analysis in [26], we form all pairs of VHH and nanobody chains and take every nonmatching pair to be putatively nonbinding (4692 pairs). We then co-fold all 69^2^ = 4761 pairs with all four methods (Promera, Boltz-2, AF-Multimer, and Protenix-1), computing the same metrics as above.

We also trained an additional confidence module in Promera specifically on the task of discriminating binding from nonbinding interfaces (Methods). Provided a model predicted structure, this module attempts to classify each contact in the prediction as correct (present in the ground truth) or incorrect (not present).

We define the *interface contact score* (iCS) as the average predicted probability across interface contacts present in the prediction and include the iCS along with ipTM and ipSAE as Promera confidence metrics.

Promera metrics again provide the strongest discrimination ability between binding and nonbinding pairs, with Promera ipSAE attaining an AUROC of 0.64 versus 0.58 for the next-best method (Protenix-v1 ipTM) (Figure 2, right). In an enrichment analysis (Figure 4), Promera iCS provides the highest enrichment for correctly matched pairs at most recall thresholds—18x at 10%, 10x at 20%, and 5x at 30%. In contrast, Boltz-2’s ipTM assigns high confidence to most pairs, regardless of correctness, and Protenix-v1’s ipTM degrades rapidly in terms of enrichment as the recall threshold increases (7.5x at 20%, 2.65x at 30%). Accordingly, when visualized as a matrix of confidence scores, only the Promera iCS is able to provide a discernible diagonal for matching VHH-antigen pairs (Figure 4, Top). We note that a threshold of AF-Multimer ipTM*>*0.7 has been used previously as a filtering threshold [28], corresponding to an ≈10x enrichment at 10% recall. Our analysis suggests that at a similar recall, Promera iCS*>* 0.8 could offer ≈20x enrichment of binders, double the expected success rate from using AF-Multimer ipTM.

**Figure 4.**
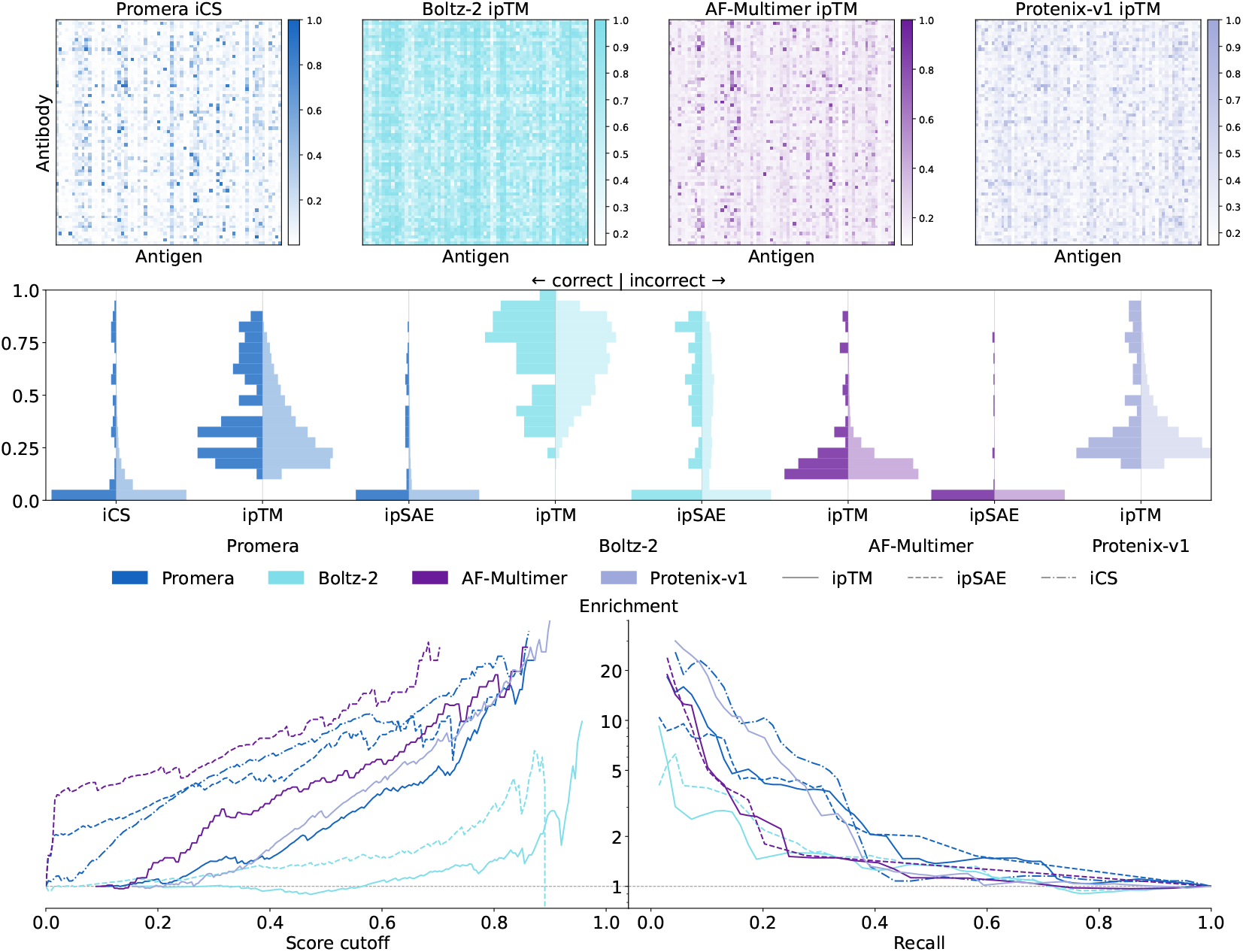
Discrimination of correct versus incorrect pairs in the shuffled nanobody-antigen dataset. (**Top**) Matrix of confidence scores for all shuffled pairs. A perfect discriminator would have ones on the diagonal and zeros elsewhere. (**Middle**) Histograms of each score for matching and non-matching shuffled pairs. (**Bottom left**) Enrichment factors computed with each scoring method, as a function of score cutoff. Note that the curves cannot be directly compared. (**Bottom right**) Enrichment-recall tradeoff curves for each scoring function. Promera’s iCS has the highest enrichment for ≈10%–35% recall. ipTM: interface predicted TMscore; ipSAE: interaction prediction score from aligned errors; iCS: interface contact score.

### Nanobody design with Promera

The promise of *de novo* binder design has driven significant interest in generative models for drug discovery [8]. Historically, binder generative models have been separately trained from co-folding models, oftentimes with different architectures or training recipes [7, 13, 27]. Here, we establish that co-folding models, with minor modifications during training, are capable binder design models. Building on the observation that co-folding models can produce structured predictions for masked sequences [11], we introduce training-time masking to further promote this behavior. At inference time, we co-fold the target sequence and a binder sequence with to-be-designed residues masked, with optional epitope, paratope, or target template conditioning. The resulting structure provides backbone coordinates for the masked residues, whose identities are then filled in with an inverse folding model. Here, we use ProteinMPNN with soluble weights, although any inverse folding model could be used.

We benchmarked the ability of Promera to design nanobodies, a class of molecules with high therapeutic value. Compared to minibinders (Figure 15), nanobodies are challenging due to the lack of secondary structure in the variable regions critical for binding and specificity. To design nanobodies with Promera, we provide the sequence of a nanobody framework (caplacizumab) with all-masked complementarity-determining regions (CDRs) of random lengths. To promote CDR contacts rather than framework contacts, we label CDR residues with a paratope flag. We selected four canonical benchmark targets from [31]— IL7Ra, InsulinR, PDGFRb, and PDL1—and sampled 10k complex structures and CDR sequences. To provide performance comparisons, we also sampled 10k designs per target with the same framework with BoltzGen using default settings. Finally, we ran 1000 trajectories (without early stopping; each producing up to 10 sequences) with mBER [28], a backprop-based protocol based on AF-Multimer. To assess the quality of all designs, we co-fold designed sequences with Promera and tabulate the best antibody-antigen iCS. Because Promera masked CDR residues are designed by ProteinMPNN, the re-folding serves as an orthogonal oracle for all design methods.

Figure 5 illustrates that Promera nanobody designs substantially outperform BoltzGen in passing iCS score thresholds (these are more stringent than ipTM thresholds of the same value; Figure 4). For IL7Ra and InsulinR, ≈40% of Promera designs pass iCS>0.6; whereas ≈80% BoltzGen samples fall below iCS<0.2. For PDGFRB, ≈75% of Promera samples pass iCS>0.6, compared to <20% for BoltzGen. Accordingly, Promera exhibits significantly higher success rates at the most stringent iCS cutoffs. We believe these higher success rates compared to the architecturally similar BoltzGen are a consequence of paratope conditioning, which avoids samples dominated by non-designable framework contacts.

**Figure 5.**
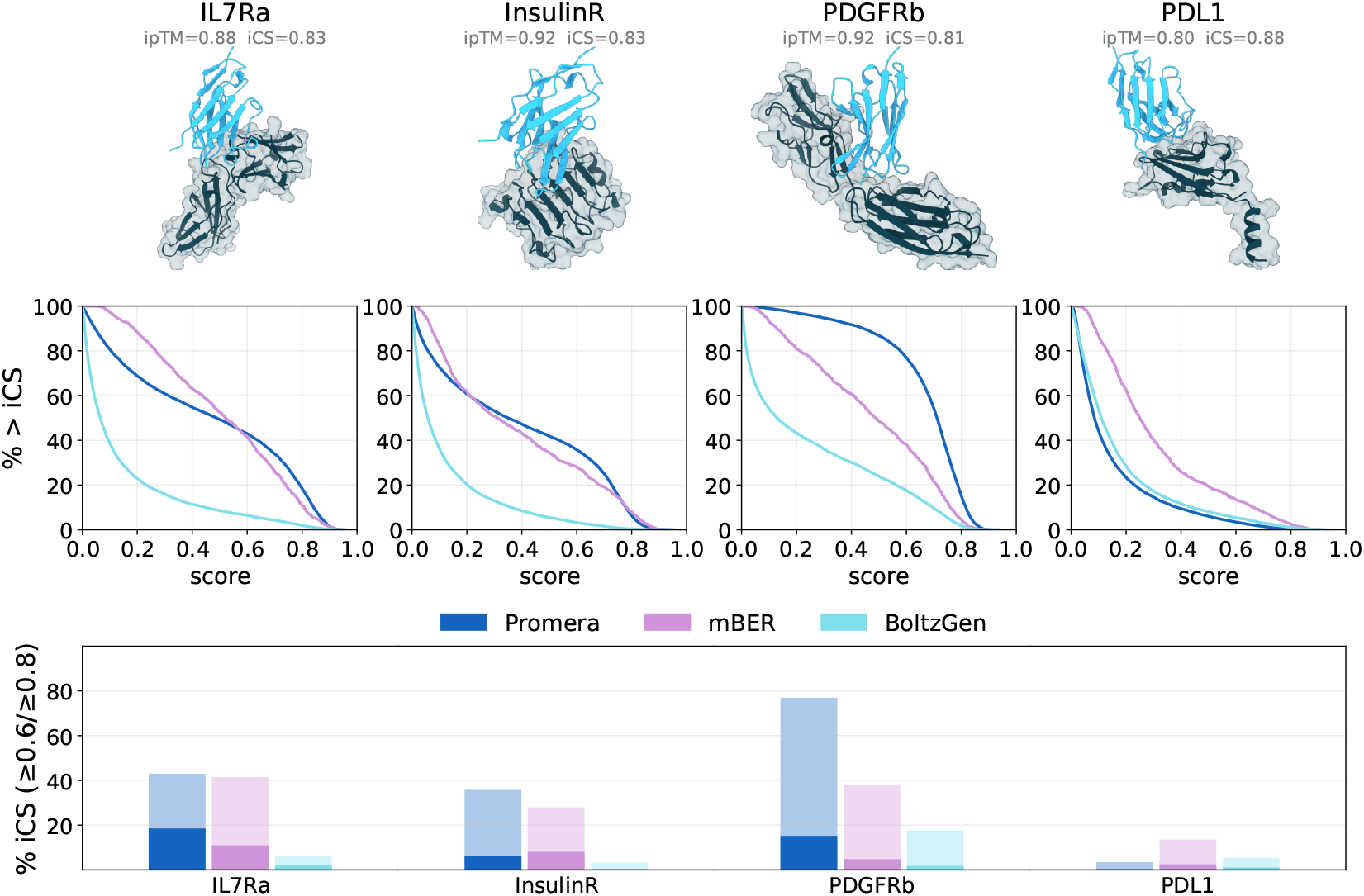
Nanobody design with Promera. compared to BoltzGen and mBER on four benchmark targets. (**Top**) Target and nanobody structures for randomly selected Promera designs with iCS > 0.8. (**Middle**) For each target and method, we plot the fraction of passing designs at each iCS threshold. (**Bottom**) Success rates of the different methods at iCS > 0.6 and iCS > 0.8 thresholds. These correspond to ≈10x and ≈20x enrichment of binders in the shuffled nanobody experiment, Figure 4. All Promera sequences are designed by ProteinMPNN from Promera structures and do not undergo any optimization for iCS. Note that Promera is not provided the target nor framework structure, unlike mBER (target structure) or BoltzGen (both).

We next compared against mBER, a nanobody-specific analog of the widely successful backprop-based method BindCraft [24]. Such backprop-based methods are generally thought to have higher per-sample success rates than generative models such as BoltzGen or potentially ours [6]. Remarkably, Promera also outperforms mBER on two out of four targets (IL7Ra, PDGFRb) and is comparable on a third (InsulinR), despite the direct optimization of co-folding confidence metrics by mBER. To our knowledge, Promera is the first diffusion-based method with *in silico* per-sample success rates comparable to backprop-based design methods.

#### Epitope targeting of the Andes hantavirus glycoprotein

A key advantage of *de novo* designed antibodies, relative to those obtained via immunization or display campaigns, is more precise control over the binding site and geometry. Such control may be desired to effect certain therapeutic mechanisms, prevent immune escape, or supplement traditional campaigns limited by immunodominance. We illustrate the epitope steering capabilities of Promera nanobody design by targeting varied epitopes of the Andes hantavirus glycoprotein. Recent hantavirus alerts in the Americas [9, 23] underscore the continued public health threat of the Andes virus. Its Gn/Gc glycoprotein complex mediates viral entry [19] and is a key target of neutralizing antibodies [18, 17], making it a compelling benchmark for controllable binder design.

To carry out an *in silico* design campaign, we selected all residues in PDB 8DBZ with high accessible surface area and cluster into eight disjoint epitopes. For each epitope, we designed and scored 10k VHHs using the same Promera protocol as described above. To score the design adherence to input conditioning, we additionally defined an *epitope engagement score* by taking the product of the scDockQ (self-consistency dockQ between the designed and refolded structure) and the fraction of labeled epitope residues in contact with the refolded VHH. For six out of eight epitopes, Promera was able to design confidently predicted VHH binders, with iCS > 0.65 and ipTM > 0.75, that at least partly contact the specified epitope residues (Figure 6). These six epitopes varied in the proportion of successful designs with high iCS and epitope engagement—an *in silico* analog of immunodominance. Nonetheless, these results demonstrate that Promera can be used to control and expand the epitope coverage of *de novo* designed libraries.

**Figure 6.**
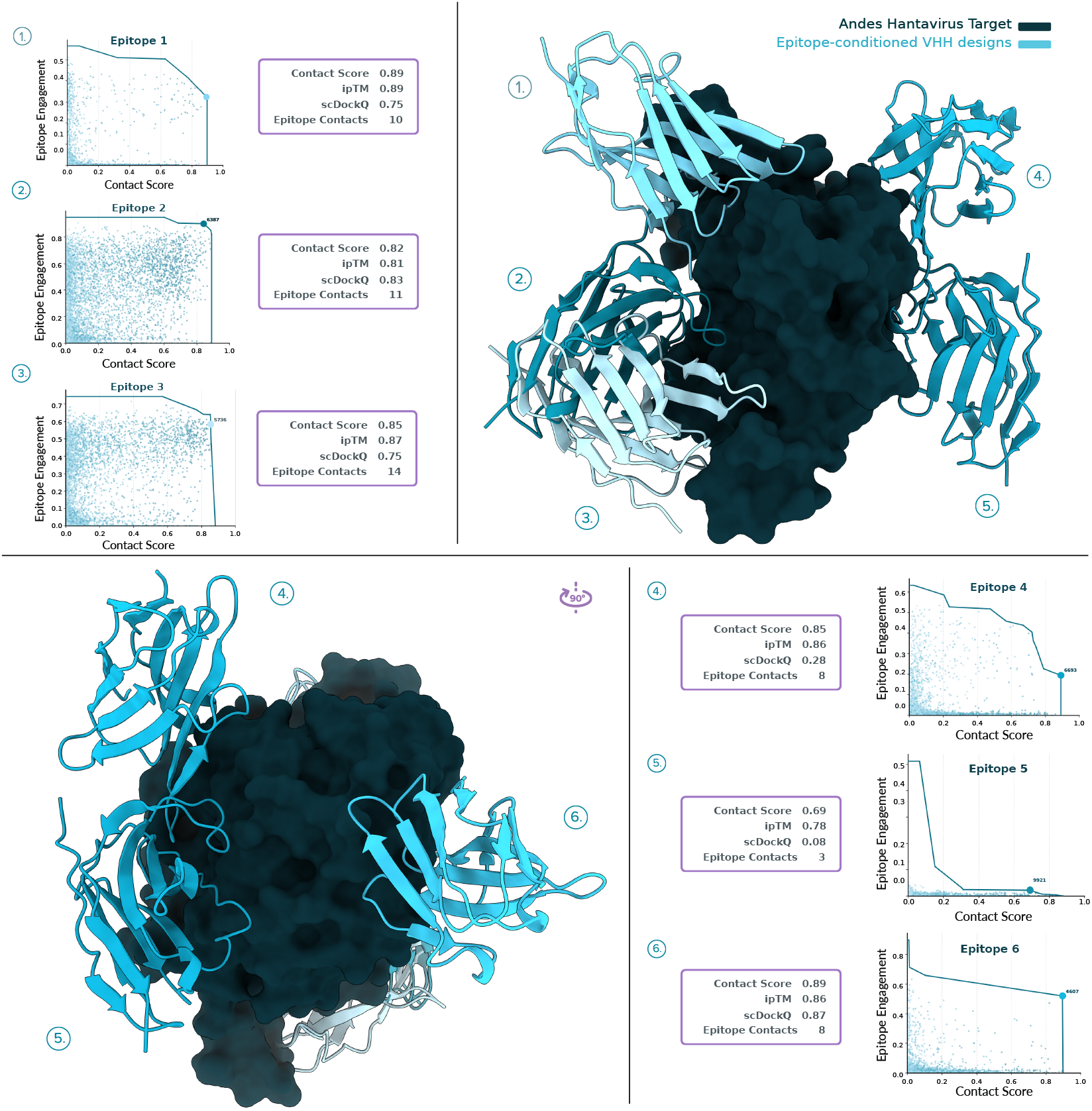
Epitope targeting of the Andes hantavirus glycoprotein. For each of six successfully targeted epitopes (out of eight), we plot 10k Promera designs, scored with iCS and epitope engagement (see text). We highlight the Pareto frontier of the tradeoff between these scores and visualize the VHH with the highest epitope engagement given iCS > 0.65. The two views (related by a 180° rotation) of the target reveal the spatial separation of the epitopes. Note that the glycosylation (if any) of the target was ignored.

#### Stabilizing the active state of the β_2_ adrenergic GPCR

Beyond binding a target epitope, *de novo* designed antibodies offer the possibility of stabilizing desired conformational states of a target. This capability would have significant value in drug discovery, as screening for functional activity cannot be easily done via display or immunization. To demonstrate this capability with Promera, we sought to design nanobodies to agonize the *β*_2_ adrenergic receptor (*β*_2_AR), a canonical member of the G-protein coupled receptor (GPCR) family. Signal transduction in *β*_2_AR activation is mediated by a crucial residue pair (I121/F282) in the transmembrane core, which switches between active and inactive conformations [14]. Remarkably, when predicting *β*_2_AR in isolation with multiple seeds, Promera produces a mixture of active and inactive states for this residue pair (Figure 7, right). Thus, we posited that a population shift towards the active state could serve as an *in silico* prediction for agonism.

**Figure 7.**
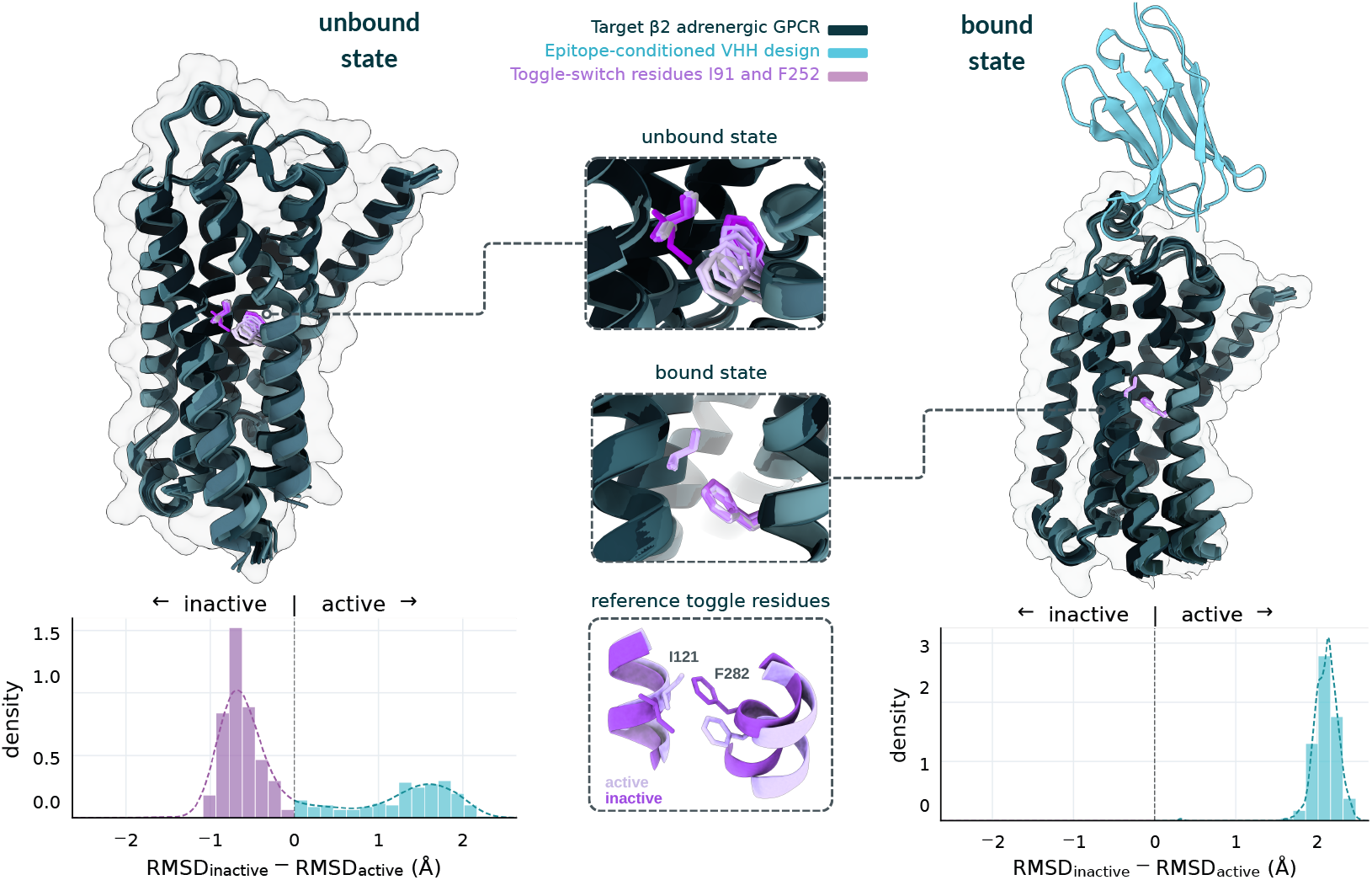
Targeting the extracellular surface of *β*_2_AR to stabilize its active state. (**Left**) Promera predictions for *β*_2_AR in isolation fluctuate between inactive- and active-like orientations of the switch residues I121 and F282, as quantified by the bimodal distribution of ΔRMSD over the toggle residues. (**Right**) Promera designed VHHs are predicted to eliminate the inactive mode and lock I121/F282 into nearly identical active orientations across the ensemble. Reference toggles are taken from PDB 2RH1/4LDL.

We conditioned Promera to generate nanobody structures interacting with the *β*_2_AR extracellular surface in its active conformation (PDB 4LDL). Out of 25k candidate designs, we filtered for iCS > 0.6 and high epitope engagement score, with 18 designs passing these filters (Appendix C). We visualize one such design in Figure 7, along with an ensemble of predicted bound structures. Remarkably, the nanobody is predicted completely abolish the inactive state in the predicted ensemble, with no occupancy even upon refolding with multiple different seeds. This suggests that orthosteric site VHH binding locks the I121/F282 connector into an activation associated state.

### Data and compute scaling explains performance across model lineages

Why do co-folding models exhibit substantial variation in accuracy, despite very similar architectures and training recipes? To seek an answer to this question, we observed that published training details varied along two main axes: training compute, including number of training steps, batch size, and diffusion multiplicity (e.g., number of diffusion training examples per pairformer forward pass); and the size of the protein monomer self-distillation set. We reasoned that co-folding performance should improve monotonically along both axes, and a simple linear fit might be able to quantitatively explain observed trends. In broader AI literature, similar fits are known as *scaling laws* and are routinely used to forecast model performance in response to increasing data and compute. In protein generative modeling, scaling laws for protein language models have been proposed to forecast performance in variant effect prediction and protein engineering [3]. A scaling law for co-folding would provide similarly concrete recommendations and cost estimates to achieve a desired level of accuracy.

To infer such a scaling law, we focus on antibody-antigen DockQ>0.23 as the main performance metric. Antibody-antigen modeling remains one of the primary limitations of co-folding models and is far from performance saturation, yet the task is of high therapeutic relevance and economic interest. Using our results and those reported in OpenFold3 [34], we calculated a normalized performance (relative to AF3) for five models: AF3, Boltz-2, Protenix-v1, Promera, and the closed-source SeedFold [37]. We then inferred the training cost from the released manuscripts and/or code repositories (details in Appendix D). The resulting analysis (Figure 8) reveals that normalized performance is positively and linearly correlated with the log of both input quantities (*p*=0.03). In particular, each 10pp improvement in normalized performance appears to roughly correspond to an ≈80% increase in self-distillation data or ≈110% increase in (diffusion) training compute. This trend suggests that observed differences between open models and their gap with AF3 could be a natural consequence of differences in training budget. Accordingly, more routine reproduction of AF3-level performance, or even improved performance, could potentially be attained via scaling.

**Figure 8.**
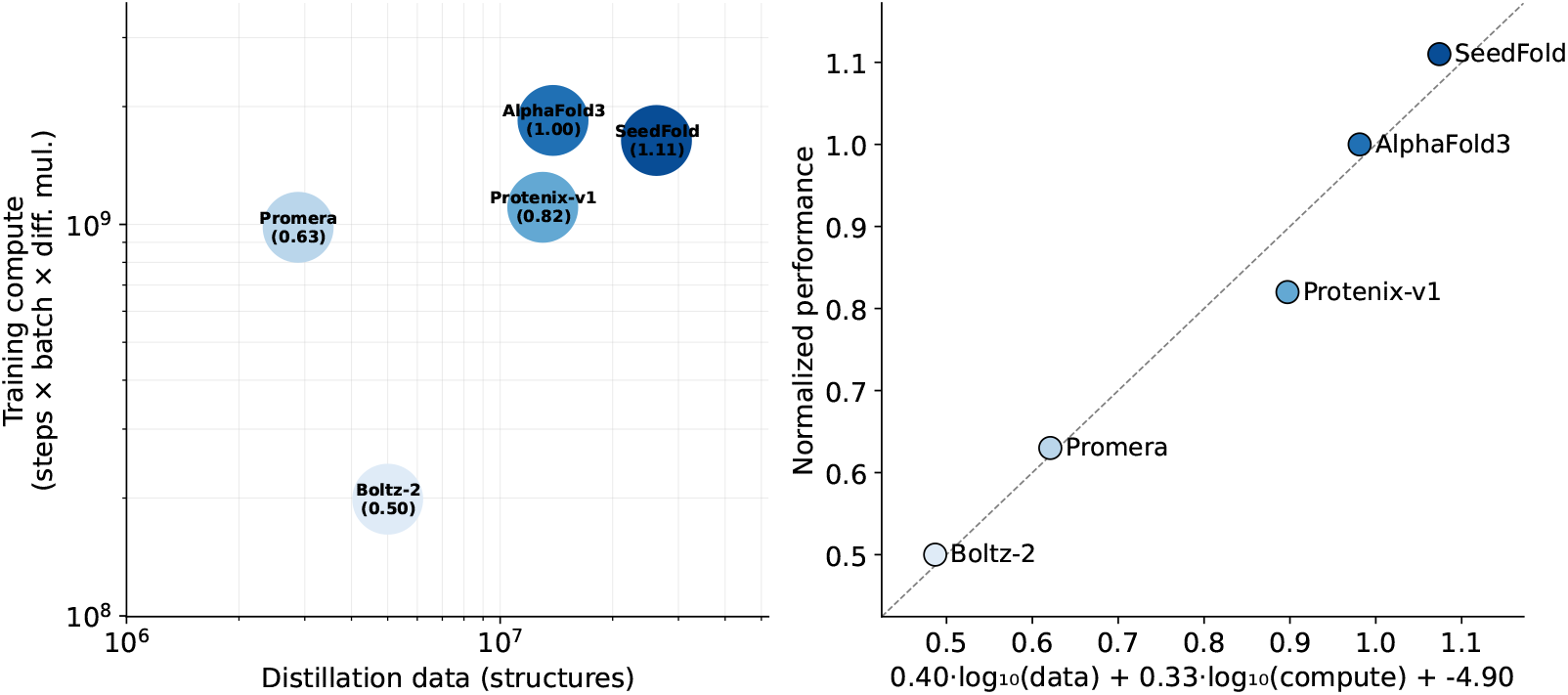
Potential scaling law for co-folding. For each method we calculate the normalized performance (relative to AlphaFold3) in terms of antibody-antigen DockQ>0.23 success rate (at 100 seeds for most methods; at 5 seeds for SeedFold). A linear function fit to the log amount of distillation data and diffusion training samples is able to model the normalized performance with high statistical significance (*R*^2^ = 0.969, *F* (2, 2) = 31.43, *p* = 0.0308).

## Discussion

We presented Promera, a unified model for both structure prediction and design. Promera achieves compelling performance across both co-folding and binder design tasks, while also addressing an overlooked but crucial step in modern design workflows: discriminating binders from nonbinders in *de novo* libraries. We provided the first systematic evaluation of binder discrimination using confidence metrics from co-folding models, which demonstrated that Promera outperforms other methods at enriching validated binders over inactive designs. We also introduce the interface contact score (iCS), a confidence metric that improves discrimination of nanobody-antigen pairs, a challenging but therapeutically valuable design category. Beyond binder discrimination, Promera supports targeted binder design through epitope and paratope conditioning, with the Andes hantavirus and *β*_2_AR case studies illustrating control over both epitope targeting and receptor state stabilization. Intriguingly, our scaling analysis suggests that co-folding accuracy improves predictability with additional training compute and self-distillation data, providing a concrete path toward stronger open models for therapeutic protein design.

## Acknowledgments

We thank Varun Ullanat, Heyuan Michael Ni, and Shorna Alam for helpful feedback and discussions. This work was supported by NIH 1R35GM141861 (to B.B.), the NSF AI Institute for Foundations of Machine Learning (IFML), the UT-Austin Center for Generative AI, and a gift from Param Hansa Philanthropies.

## A Architecture and Training Details

### Architecture

Our architecture closely follows that of AlphaFold3 and is based on the Boltz-1 implementation. However, we make the following deviations:

- Although backprop-based design protocols are popular for AlphaFold2-style models, their transfer to all-atom co-folding models [10] is hindered by the dual-track embedding of amino acid identities: via the one-hot embedding (which can be relaxed for backpropagation) and a gathered atomic embedding (which cannot). Our solution is to mask out the gathered atomic embedding for standard tokens.
- To prevent leakage of the atomic embeddings to nonstandard tokens via windowed attention, we remove the input embedding transformer prior to the pairformer.
- To facilitate design use cases, we provide an epitope conditioning flag for tokens in contact with another chain.
- Sparse pairs of intertoken distances can be provided via a partial distogram for target conformation conditioning.
- We identify a potential ambiguity in the featurization scheme of small molecules in existing co-folding models. Specifically, the standard practice of providing only the connectivity graph, formal charge, and 3D coordinates of heavy atoms can sometimes fail to discriminate between similar molecules (Figure 9). We patch this ambiguity by providing the total hydrogen coordination count for each heavy atom.

**Figure 9.**
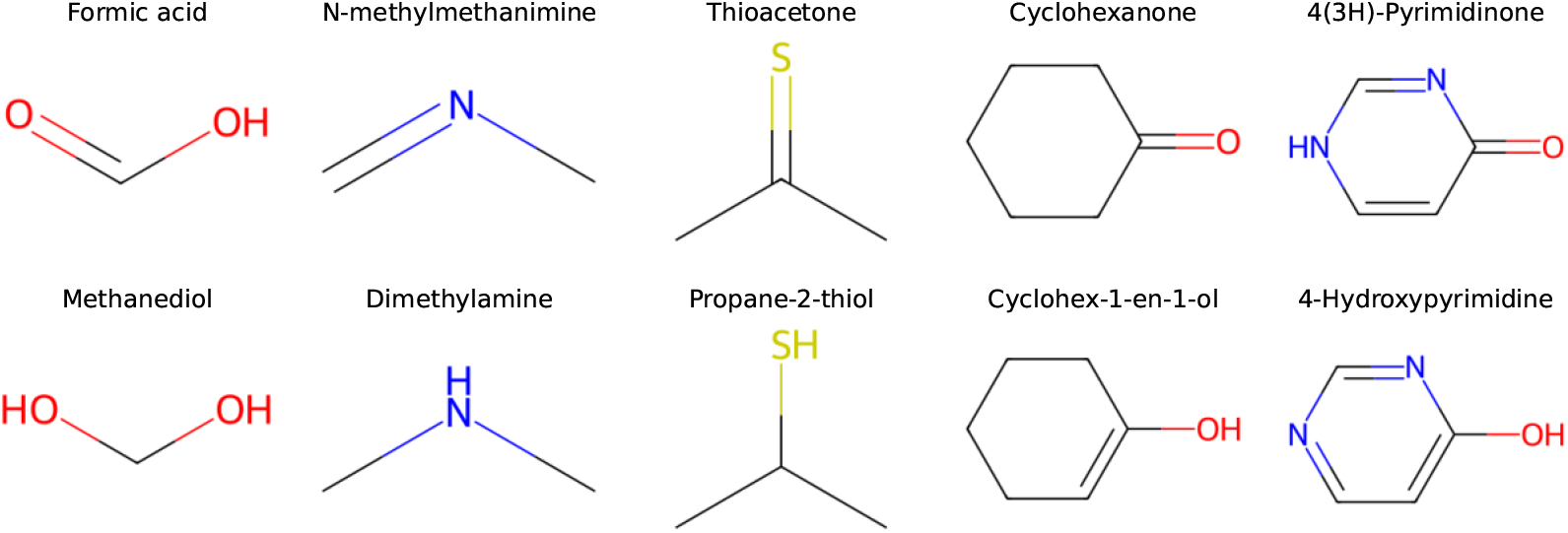
Molecule pairs which have potentially ambiguous representations under standard featurization schemes for co-folding models (heavy-atom coordinates, connectivity graph, and formal charge). In the first three examples, the molecules differ in hybridization and so in principle can be distinguished by bond geometry; however, this input may be unreliable for input conformers.

### Training

As is standard practice, we supplement PDB training with a self-distillation monomer set. To curate this distillation set, we start with the FoldSeek clustering of Mgnify [36] to identify clusters that do not have any AlphaFoldDB members. We then filter cluster representatives for length ≥ 200 and re-predict structures using ColabFold [22], resulting in 1.13M structures. We supplement these structures with the 3.35M AFDB members with length ≥ 200 whose MSAs are in OpenProteinSet [2]. At training time, we sample from the PDB, Mgnify, and AFDB with a 50%/35%/15% ratio. To align the model training with backbone generation for design, we replace a fraction of standard amino acids with `UNK` tokens and remove their sidechain atoms in 5% of batches. Similarly, we provide epitope conditioning and partial distogram conditioning each in 5% of batches. Training proceeds in three stages: 80k steps with crop size 512, 20k steps with crop size 640, and finally 20k steps with crop size 768.

**Figure 10.**
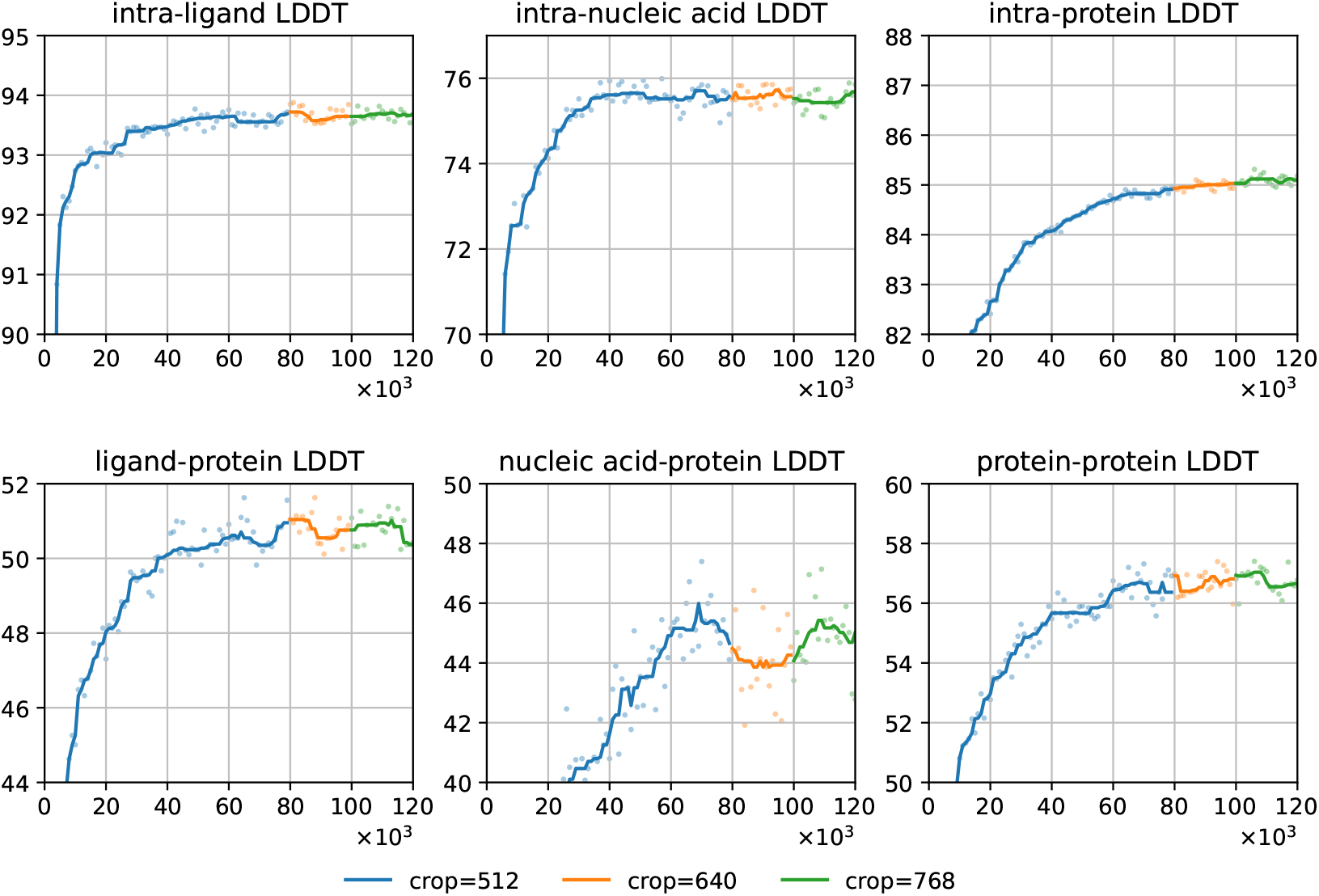
Validation accuracy, stratified by category, across three training stages tracked on selected 2024 PDB complexes. Lines are smoothed with a median filter of window size 9.

## B Evaluation Details

### PDB test set

Our test set starts with the previously described PXM-25 set [20], which comprises 2359 complexes. We then filtered the set to ensure consistent evaluations across methods (i.e., removing input modalities not supported by all methods) and to reduce evaluation latency. In particular, we retained biological assemblies satisfying all of the following conditions: (1) no branched entities, (2) no covalently bound ligands, (3) 100 or fewer chain symmetries, (4) no ligand with more than 24 symmetries, and (5) 2048 or fewer tokens. The final test set comprised 1605 complexes. Following the protocols in PXMeter [20], we extract low-homology chains and interfaces, assign them to clusters, and report cluster means.

### Inference

All methods are provided MSAs from a local `mmseqs` webserver; however, unlike previous co-folding models, Promera does not use any pre-paired MSAs (`mmseqs pairaln`) for inference, relying entirely on local pairing of filtered single-chain MSAs. Although this theoretically places our model at a disadvantage, we aim to establish a precedent for lower-overhead inference workflows. For the recent PDB set, we draw 25 samples (5 seeds *×* 5 diffusion samples) with 10 recycling steps, and select the best sample according to 0.8*×* ipTM + 0.2*×* pTM. For recent anbitody-antigen complexes, we extract and co-fold the antigen and antibody chains only. We evaluate all methods with 100 seeds, 10 recycles, and 5 samples per seed, rank samples by complex ipTM, and calculate the average dockQ across antibody-antigen interfaces. For filtering *de novo* binders (from the Adaptyv competitions and designed), an MSA is provided for the target but not the design, inference is run with 4 recycles; and (for all methods except AF-M) we draw 5 diffusion samples. For the shuffled nanobody evaluation, we provide MSAs for both chains, but omit paired MSAs. All confidence metrics are taken as the maximum over the 5 samples.

### Stratified confidence correlations

To assess the utility of co-folding confidence metrics, we report several different pearson correlation coefficients. Let *y*_*ijk*_ be the accuracy metric (LDDT or dockQ) of complex *i*, seed *j*, and sample *k*, and let *ŷ*_*ijk*_ be the corresponding confidence prediction. We then report

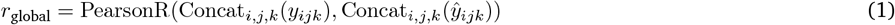

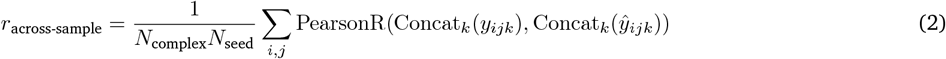

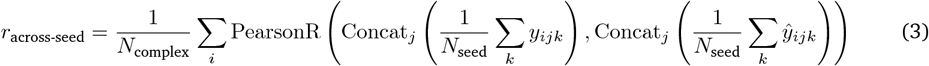

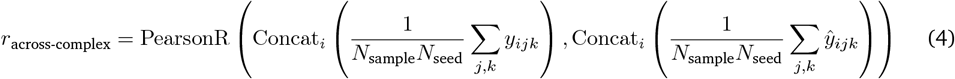

## C Additional Results

### C.1 Confidence module quality

The ability of confidence modules to correctly identify the best sample among multiple predictions is essential for inference scaling and self-distillation. We explicitly benchmark the confidence modules of Promera, Boltz-2, OF3, and Protenix-v1 using predictions on the recent PDB test set and the antibody-antigen set (Figure 11). By stratifying the correlation coefficients, we disentangle the ability to discriminate multiple predictions for the same complex versus ranking complexes by difficulty (Appendix B). While methods generally have moderate ability (*r* ≥ 0.4) to report the difficulty of a complex, they have weaker ability (*r* ≤ 0.2) to discriminate between predictions within a complex. Nevertheless, differences between methods can be meaningful: on the antibody-antigen dataset, Protenix-v1 has the highest within-complex discrimination ability, consistent with its improved inference scaling behavior. The generally higher correlation coefficient across seeds versus across samples is consistent with the observation that increasing diffusion samples is not effective at improving predictions [1]. We anticipate that careful consideration of confidence module design and training will be important to further advances in co-folding performance.

**Figure 11.**
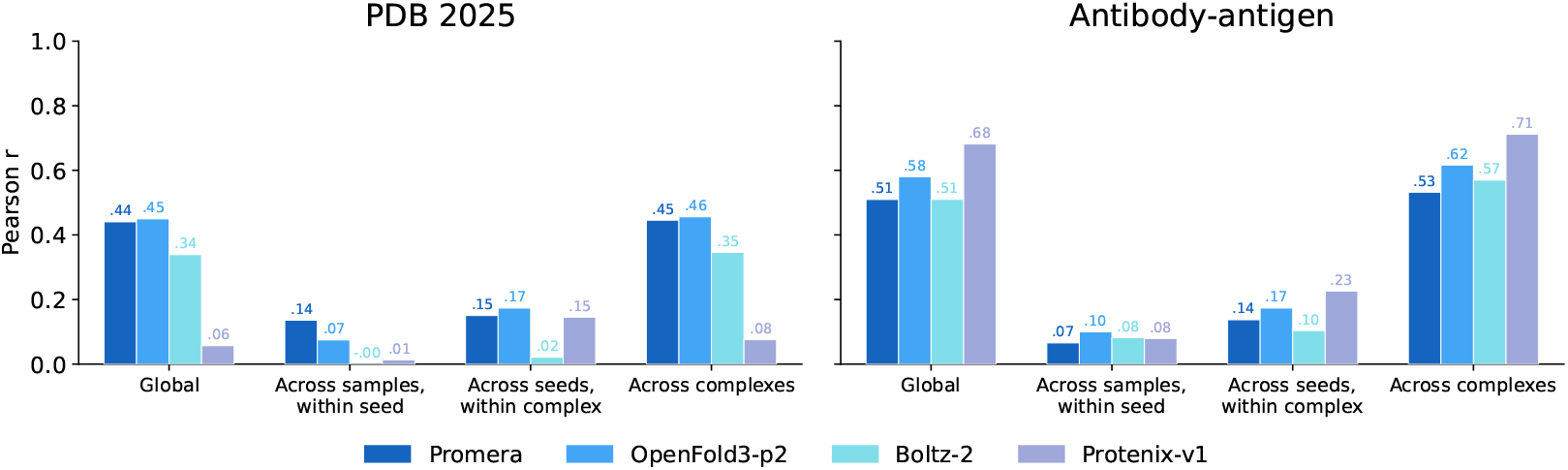
Analysis of confidence model quality. For each method, we compute the Pearson correlation between ranking score (0.8*×* ipTM + 0.2*×* pTM) and complex LDDT (PDB 2025) or ipTM and dockQ (antibody-antigen). We report four coefficients to quantify the ability to discriminate (1) global quality, between different samples within a seed, (3) between different seeds for the same complex, and (4) across complexes. The global variance is dominated by the variance across complexes; however the ability to discriminate between samples and seeds is more relevant for inference scaling and self-distillation.

### C.2 Inference scaling for antibody-antigen and protein-ligand co-folding

**Figure 12.**
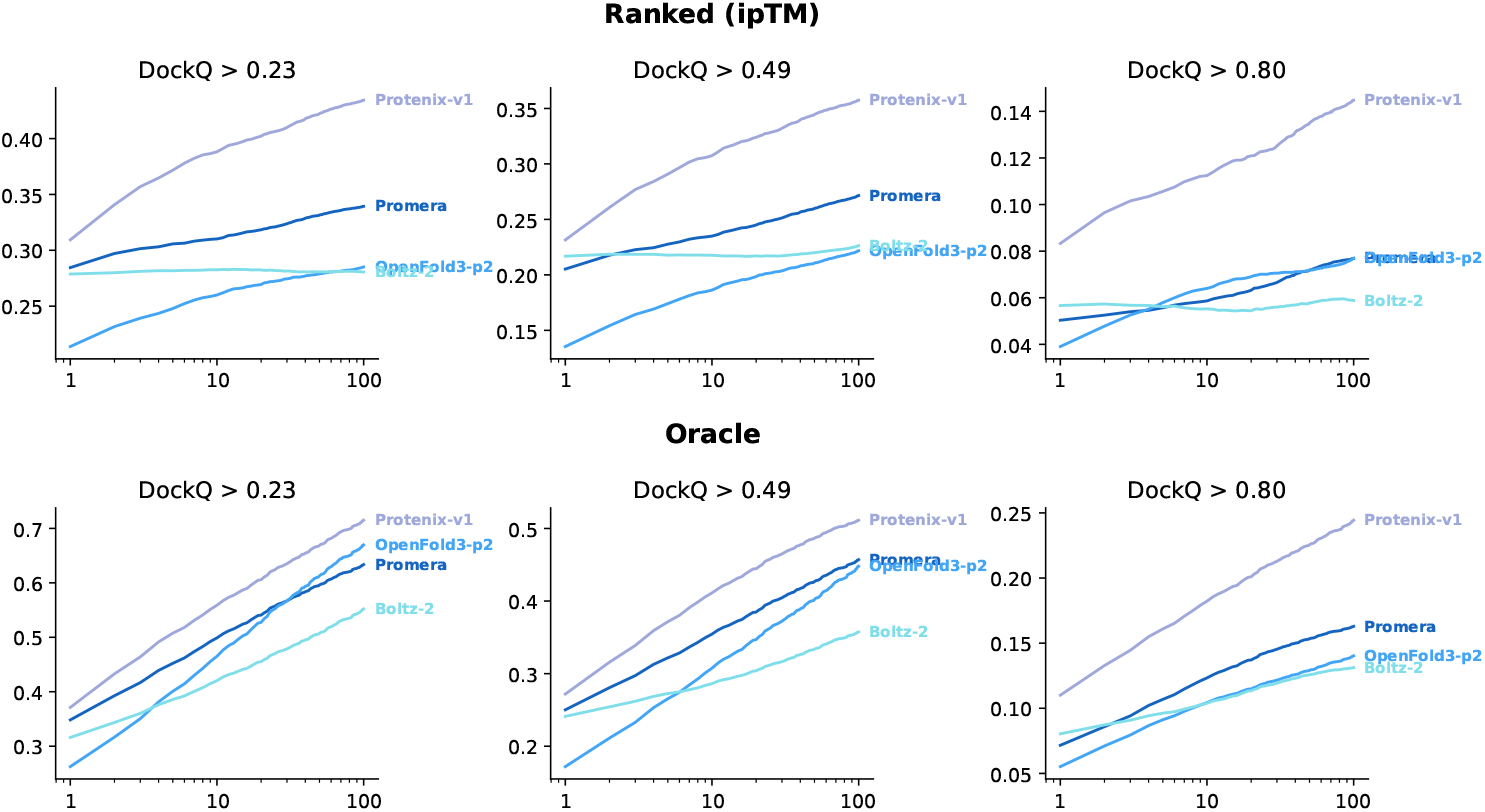
Antibody-antigen co-folding accuracy as a function of the number of inference seeds, where the best sample is selected based on ipTM (top row) or oracle DockQ (bottom row). Out of a total seed pool of 100, we simulate the accuracy for 1-99 seeds by averaging over 200 trials for each. Note that Promera is not provided pre-paired MSAs, unlike the other methods.

**Figure 13.**
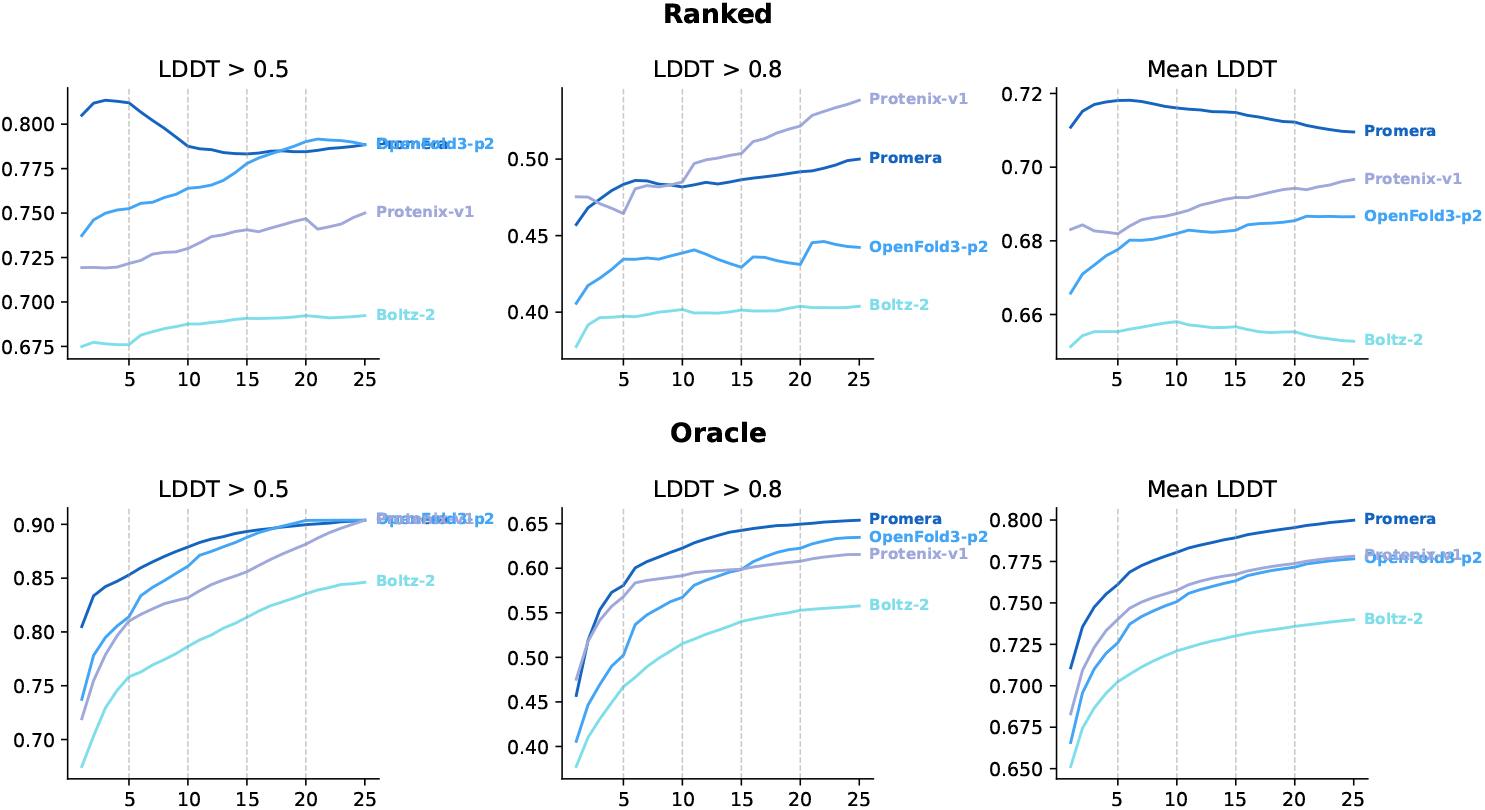
Protein-ligand co-folding accuracy as a function of the number of samples, grouped by seed. Out of the total pool of 5 seeds *×* 5 samples, we simulate the accuracy at lower sample counts by averaging over 200 repermutations (of seeds and of samples within seeds) for each complex. The best sample is selected by the overall complex confidence (top row) or by oracle protein-ligand LDDT (bottom row). Results are reported for 187 interfaces grouped into 52 clusters from the 2025 PDB test set.

### C.3 EGFR binder competition

**Figure 14.**
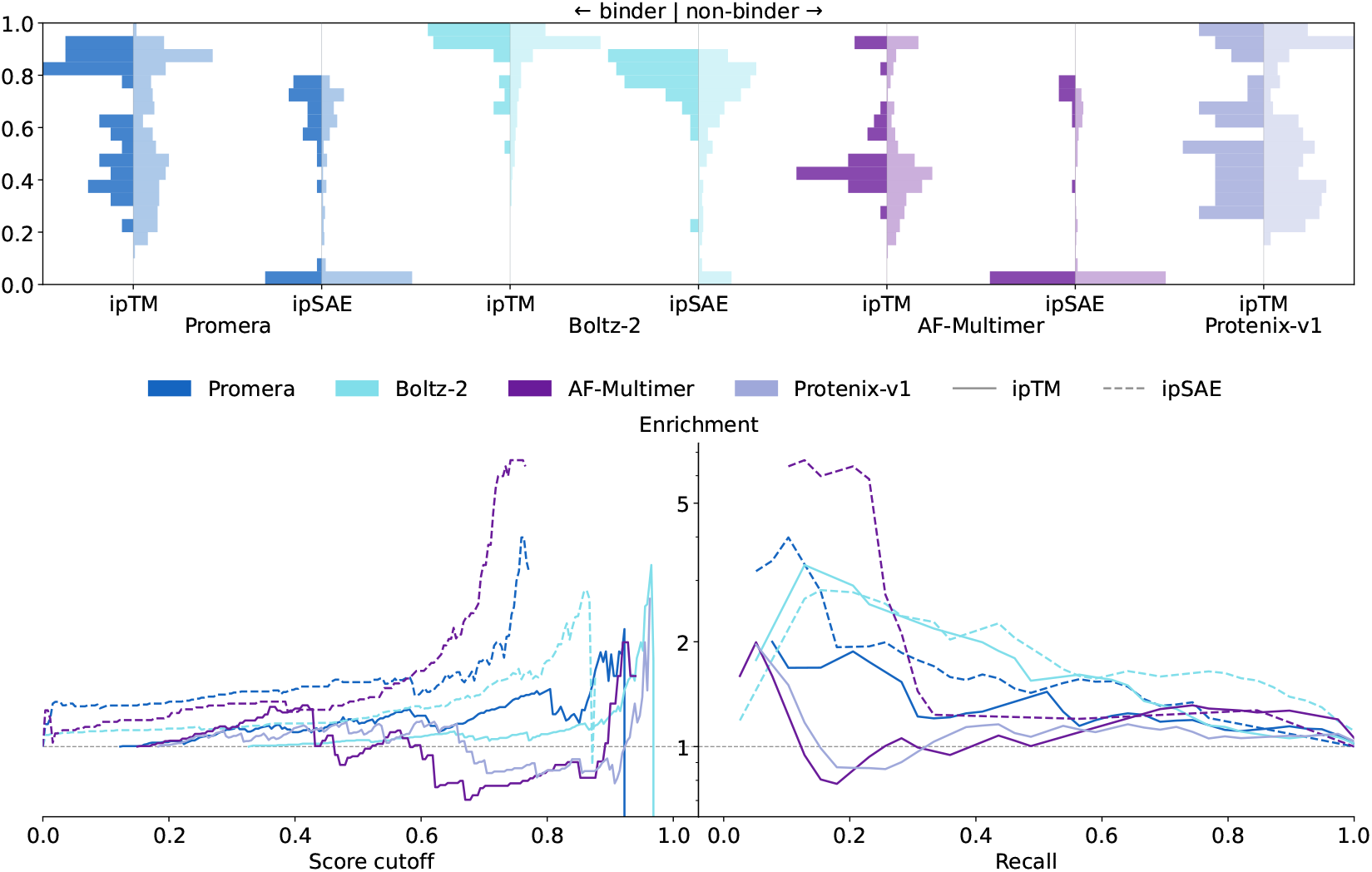
Discrimination of binders in the EGFR protein design competition with co-folding confidence metrics. (**Top**) Histograms of each score for binders and non-binders. (**Bottom left**) Enrichment factors computed with each scoring method, as a function of score cutoff. (**Bottom right**) Enrichment-recall tradeoff curves for each scoring function. ipTM: interface predicted TMscore; ipSAE: interaction prediction score from aligned errors.

### C.4 Promera-designed minibinders

We benchmark minibinder generation with Promera on four canonical benchmark targets from [31]— IL7Ra, InsulinR, PDGFRb, and PDL1. For each target, we predict 10k structures with a masked binder sequence (length 40–120), and design one sequence per structure with ProteinMPNN. For comparison, we also draw 10k designs from BoltzGen [27] using default settings, which also provide the target structure. To evaluate these designs, we co-fold all designs with Promera and 4 recycles, providing an MSA only for the target, and tabulate the maximum ipTM across five samples. Because all Promera designed sequences are sampled from ProteinMPNN, the re-folding serves as an orthogonal oracle for both design methods.

We plot the fraction of designs passing each threshold of Promera ipTM for all targets and both methods (Figure 15). We find that compared to BoltzGen, Promera designs have a generally comparable distribution of ipTM (median ipTM higher on two out of four targets), but somewhat narrower range. Using an intermediate threshold of ipTM*>*0.7, Promera designs are more successful on two out of the four benchmark targets (IL4Ra and InsulinR). We note that since BoltzGen by default uses the PDB structure of the target, these results may not represent a fully controlled comparison.

**Figure 15.**
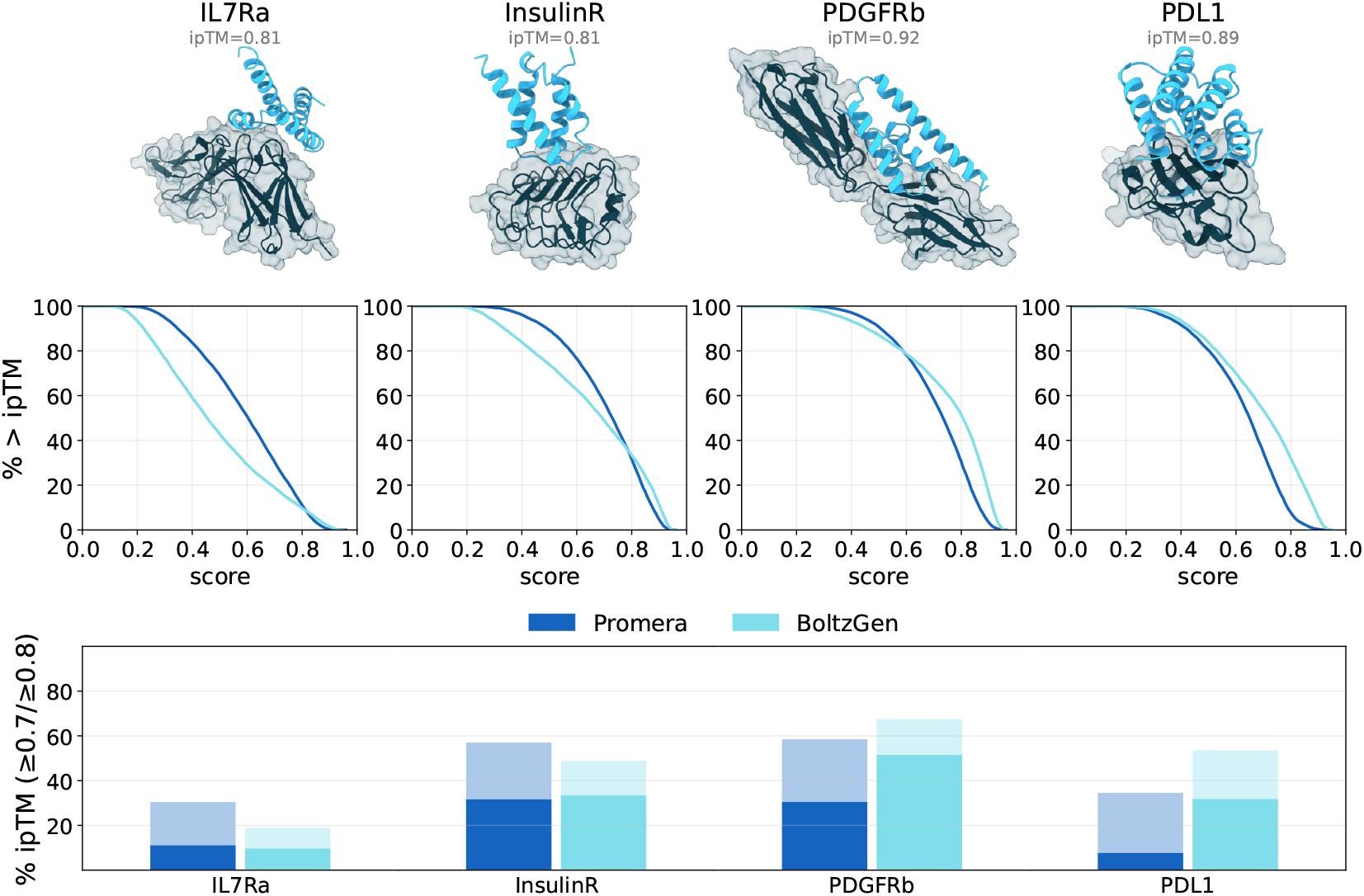
Minibinder design with Promera compared to BoltzGen on four benchmark targets. (**Top**) Target and nanobody structures for randomly selected Promera designs with ipTM > 0.8. (**Middle**) For each target and method, we plot the fraction of passing designs at each ipTM threshold. (**Bottom**) Success rates at ipTM > 0.7 and ipTM > 0.8 thresholds.

### C.5 *β*_2_AR designs

**Figure 16.**
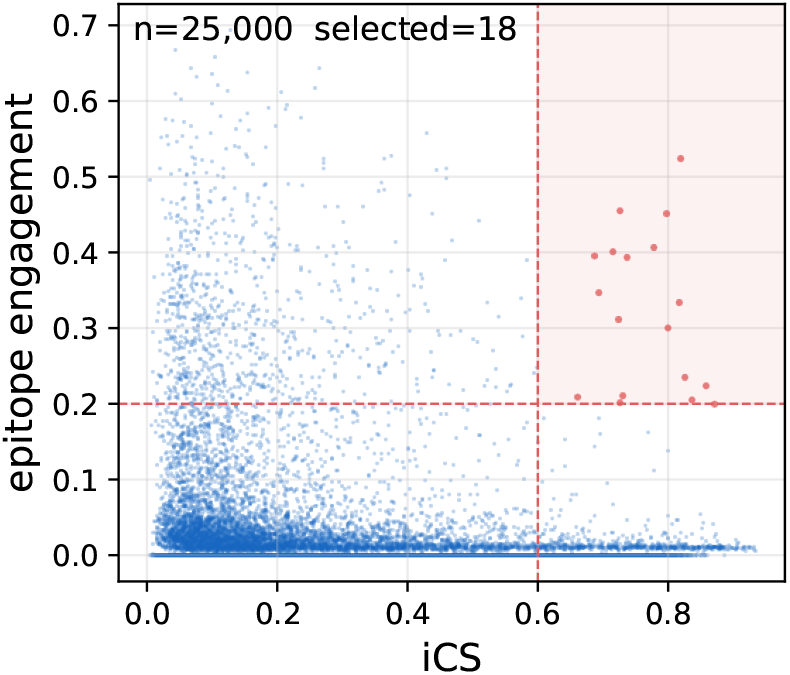
All generated designs targeting the orthosteric site of *β*_2_AR. Epitope engagement is the product of scDockQ and the fraction of specified epitope residues that are in contact with the refolded VHH.

## D Scaling Law Estimation

We estimate the total training inputs of Boltz-2, Protenix-v1, SeedFold, and AlphaFold3 from their published reports and/or config files in their respective codebases. Note that the diffusion multiplicity is sometimes decreased over the course of training; we take the value reported for the majority of training. If there are multiple protein monomer distillation sets, we compute the effective single-dataset size based on a uniform sampler with the same entropy.

**Table 1:** Estimation of compute and data inputs for the co-folding scaling law.

| Model | Parameter | Value | Source |
| --- | --- | --- | --- |
| AlphaFold3 | Diffusion mul. | 48 | Manuscript |
|  | Batch size | 256 | Manuscript |
|  | Training steps | 150k | Extended Data Fig. 2 |
|  | Distillation structures | 13.85M | Manuscript (1% 28M + 99% 13M) |
| Protenix-v1 | Diffusion mul. | 48 | https://github.com/bytedance/Protenix/blob/main/configs/configs_base.py |
|  | Batch size | 256 | Manuscript |
|  | Training steps | 90k | Protenix-v0.5 manuscript [29] |
|  | Distillation structures | 13M | Manuscript |
| Boltz-2 | Diffusion mul. | 16 | https://github.com/jwohlwend/boltz/blob/main/scripts/train/configs/full.yaml |
|  | Batch size | 128 | https://github.com/jwohlwend/boltz/blob/main/scripts/train/configs/full.yaml |
|  | Training steps | 97k | Manuscript |
|  | Distillation structures | 5M | Manuscript |
| SeedFold | Diffusion mul. | 64 | Manuscript |
|  | Batch size | 256 | Manuscript |
|  | Training steps | 100k | Manuscript |
|  | Distillation structures | 26.2M | Manuscript (16% 3.3M + 84% 23M) |

